# Structural mechanisms of allosteric regulation in the human *cis*-prenyltransferase complex

**DOI:** 10.1101/2025.05.09.652838

**Authors:** Moshe Giladi, Shiri Kredi, Carlo Guardiani, Lyr Aviram, Pavla Vaňková, Petr Man, Alberto Giacomello, Yoni Haitin

## Abstract

Human cis-prenyltransferase (h*cis*-PT) synthesizes long-chain isoprenoids essential for N-linked protein glycosylation. This heteromeric complex comprises the catalytic subunit DHDDS and the regulatory Nogo-B receptor (NgBR). Although NgBR dramatically enhances DHDDS activity, the molecular basis for this allosteric regulation remains unclear. Here, we combined crystallography, hydrogen-deuterium exchange mass spectrometry (HDX-MS), molecular dynamics simulations, and network analysis to uncover the structural dynamics and communication pathways within h*cis*-PT. By solving the apo structure of h*cis*-PT, we reveal only a localized flexibility at the active site and the NgBR C-terminus. However, HDX-MS demonstrated widespread substrate-induced stabilization, particularly at the NgBR βD–βE loop, highlighting it as an allosteric hub. Functional mutagenesis scanning identified NgBR^S249^ as critical for enzymatic activity, independent of structural perturbations. Network analysis of MD simulations pinpointed this residue as a central node in inter-subunit communication, with perturbations disrupting downstream allosteric pathways, altering enzymatic activity. Our findings reveal a dynamic regulatory network centered at the inter-subunit interface, wherein specific NgBR residues modulate DHDDS activity through allosteric signaling. This work elucidates a conserved mechanism of subunit coordination in long-chain cis-prenyltransferases and suggests novel avenues for therapeutic targeting of h*cis*-PT-related disorders.

## Introduction

Isoprenoids are a vast group of chemically diverse compounds essential for a myriad of cellular processes in all life forms^1^. These compounds serve as visual pigments, hormones, membrane constituents, and post-translational protein modification moieties and participate in bacterial cell wall synthesis, among other roles^2^. The prenyltransferase enzyme superfamily is responsible for the synthesis of isoprenoids by catalyzing the condensation reaction of a common precursor, isopentenyl diphosphate (IPP, C_5_), onto allylic diphosphate primers of varying lengths^3–5^. This superfamily can be subdivided into the structurally and catalytically distinct *cis*- and *trans*-prenyltransferase branches, according to the stereoisomerism of the double bond formed^1,3–6^. The *cis*-prenyltransferase branch is further classified by the product chain length^1^. While enzymes synthesizing short- and medium-chain products (C_10_-C_60_) form homodimeric complexes, long-chain isoprenoids are formed by heteromeric complexes^1,7,8^.

The human *cis*-prenyltransferase complex (h*cis*-PT), belonging to the long-chain class, is a key cellular enzyme associated with the endoplasmic reticulum (ER) membrane and ubiquitously expressed in every cell type^7,9^. Like other long-chain *cis*-prenyltransferases, h*cis*-PT is a heteromer consisting of the catalytic dehydrodolichyl diphosphate synthase (DHDDS) and the auxiliary Nogo-B receptor (NgBR) subunits^7^. Catalyzing the elongation of farnesyl diphosphate (FPP, C_15_) via multiple condensations with IPP, h*cis*-PT synthesizes DHDD (C_85-100_)^7^. DHDD is the precursor for dolichol-phosphate (Dol-P), the glycosyl carrier molecule necessary for N-linked protein glycosylation (Fig. 1)^10,11^. As this modification is essential for protein folding, oligomerization, quality control, sorting, and transport^10^, even slight variations in protein glycosylation can markedly affect protein structure and function^12–14^, emphasizing the significance of h*cis*-PT to numerous cellular processes.

**Figure 1.**
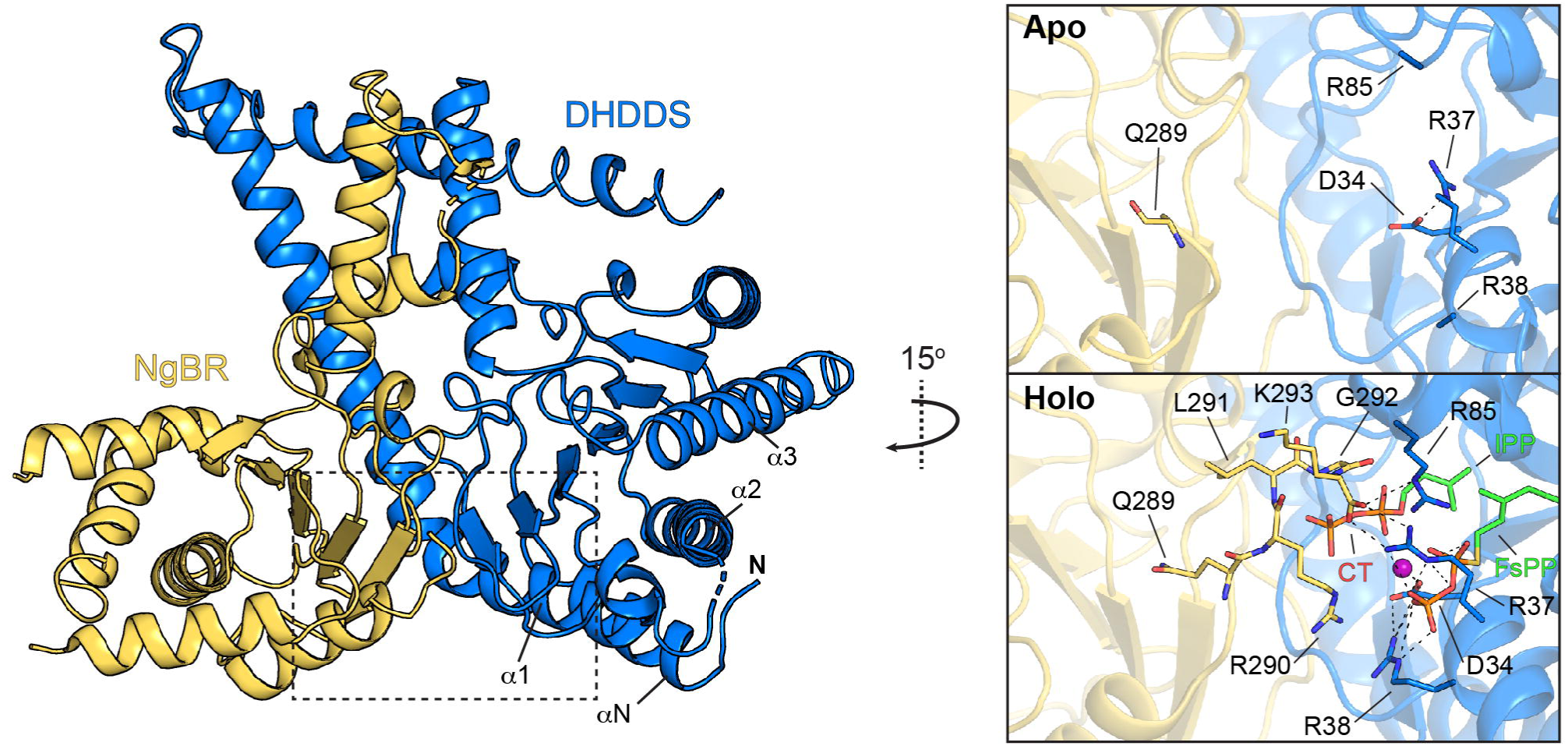
Crystal structure of h*cis*-PT in the apo state. Cartoon representation of the crystallographic asymmetric unit, composed of a DHDDS (blue) and NgBR (yellow) heterodimer. Active site secondary structure elements of interest are labeled. The dashed rectangle frames the NgBR C-terminus and the DHDDS diphosphate binding region, shown in the zoom perspective on the right. The holo structure, with bound FsPP and IPP (PDB 7PAX), is shown for comparison. Note that in the apo state, the NgBR C-terminus as well as the side chains of the catalytic residues R38 and R85 could not be resolved, due to enhanced flexibility. In addition, a novel salt bridge between R37 and D34 of DHDDS is formed.

In line with the cellular significance of h*cis*-PT, mutations in both subunits were implicated in human diseases involving practically every organ system^15^. Intriguingly, the retina and the brain are particularly susceptible to the deleterious effects of DHDDS mutations, with phenotypes ranging from non-syndromic retinitis pigmentosa (RP)^16,17^ – a progressive degeneration of photoreceptors leading to vision loss – through developmental epileptic encephalopathies (DEE)^18–22^ – severe neurological conditions characterized by frequent seizures and developmental delays. However, a multi-organ and fatal congenital disorder of glycosylation (CDG) was also reported due to mutations occurring in either subunit of the h*cis*-PT complex^22,23^. While altered Dol-P levels are expected to result in defective cellular glycosylation, direct evidence for such a pathophysiological linkage awaits further investigations^24,25^.

The heteromeric assembly of DHDDS and NgBR subunits serves a vital functional purpose, possibly contributing to catalytic activity and/or product chain-length determination^1^. Indeed, comparing the enzymatic activity of purified homodimeric DHDDS to that of the heteromeric h*cis*-PT revealed that NgBR confers an activity increase of ∼400-fold^26–30^. In addition, NgBR was shown to be necessary but insufficient for dolichol synthesis^7^. However, the structural mechanism underlying this effect remains elusive. Interestingly, the distal C-terminus of NgBR encompasses a conserved ^290^RxG^292^ motif, in which a lethal CDG mutation (R290H) was previously characterized in a family of Roma origin^22^. The structures of h*cis*-PT revealed that this NgBR region traverses the inter-subunit interface, forming an integral determinant of the DHDDS active site architecture. However, while NgBR^R290^ interacts with the substrate and catalytic residues, *in vitro* enzyme kinetics analysis of the NgBR^R290H^ mutation resulted in a mere 30% decrease in enzymatic activity. Similar reductions in catalytic activity were obtained in other mutants of the NgBR C-terminus^31^, pointing towards the involvement of a complex allosteric network governing the functional regulation of DHDDS catalytic activity by NgBR.

Despite the availability of high-resolution structures of the h*cis*-PT complex along its catalytic cycle^26–28^, structural comparisons did not reveal apparent conformational changes in NgBR that would explain its marked impact on DHDDS function. Here, therefore, we first explored the structural dynamics of the complex in different active site occupancy states by obtaining the crystallographic structure of h*cis*-PT in its apo state and using hydrogen-deuterium exchange mass spectrometry (HDX-MS)^32^ to pinpoint NgBR regions potentially involved in inter-subunit communication. Next, we delineated the communication paths traversing the complex by combining high-resolution structural analysis, molecular dynamics (MD) simulations, and network analysis^33^. Finally, we experimentally validated these allosteric networks by examining the functional ramifications of systematic perturbations of key residues along the paths. Together, our results identify the inter-subunit interface as a node for allosteric information transfer between the NgBR and DHDDS subunits, providing molecular insights into the modulatory role of auxiliary subunits in heteromeric *cis*-prenyltransferases.

## Results

### Structure of the hcis-PT complex in the apo-form

Recent structural studies of the h*cis*-PT complex resolved the active site conformation in various substrate- and product-bound states^26–28^. These structures are very similar, sharing a high degree of conformational restraint. To explore whether substantial conformational rearrangements occur upon ligand binding, we determined the 2.9 Å resolution crystal structure of h*cis*-PT in its apo-form (Fig. 1). Focusing on the active site, several regions exhibit increased conformational flexibility. Specifically, the α1-α2 linker, immediately downstream of a retinitis pigmentosa (RP59) mutation position DHDDS^K42E 16,17^, is destabilized, rendering residues ^44^QV^45^ unresolved. Moreover, the side chains of the catalytic residues DHDDS^R38^ and DHDDS^R85^, which are directly involved in coordinating the diphosphate groups, are unresolved. A novel salt bridge is observed between DHDDS^R37^, which coordinates the diphosphate group of the elongating product, and DHDDS^D34^, crucial for magnesium binding and catalytic activity^26^. Moreover, the NgBR C-terminus was not resolved beyond residue NgBR^289^, indicating conformational heterogeneity of the ^290^RxG^292^ motif. Notably, in the substrate-bound conformation, salt bridges between both DHDDS^R37^ and DHDDS^R85^ of DHDDS and the terminal carboxylate group of NgBR stabilize this region^27^. Therefore, in the absence of a ligand, changes in local electrostatics induce swapping between the DHDDS^R37^ salt-bridging pair alternates. This may explain the detrimental effect of the DEE mutation DHDDS^R37H^, which is expected to destabilize both the apo and the holo states^18^. Together, the structure of the apo-form exposes a ligand-dependent reorganization of molecular determinants directly involved in diphosphate group binding without additional global conformational changes.

### HDX-MS exposes the inter-subunit interface as a hub for allosteric communications

Given the strong regulatory effect conferred by NgBR on DHDDS activity, we sought to explore additional ligand-elicited conformational changes extending beyond the active site boundaries. As enzymatic catalysis is an inherently dynamic process^34^, some of these intermediate states may represent a rare occurrence that is not readily accessible to high-resolution crystallographic structural analysis. In HDX-MS, the exchange of backbone amide hydrogens with deuterium, reflecting the secondary structure and solvent accessibility, is monitored over time^35^. We compared the HDX-MS profiles of h*cis*-PT in its apo, FPP (substrate-bound), and GGPP (first condensation product; C_20_) states. Consistent with the crystallographic structures, binding of either FPP or GGPP resulted in decreased HDX throughout the active site, most pronounced in the diphosphate binding region encompassing residues DHDDS^D34^, DHDDS^R37^, and DHDDS^R85^ (Fig. 2). Conversely, the distal hydrophobic outlet of the active site (DHDDS^αN-βA^, DHDDS^α3-βC^) exhibited increased HDX following ligand binding. This is consistent with its previously suggested role in product egress into the membrane^27^, possibly reflecting the priming of this region for product release. Strikingly, upon occupying the active site of DHDDS by either FPP or GGPP, we observed diffused reduced conformational dynamics throughout NgBR (Fig. 2). The stabilizing effect was most pronounced at the NgBR^βD-βE^ loop (residues 244-262), situated at the interface with DHDDS. Therefore, HDX-MS exposes diffused ligand-mediated conformational dynamics spanning large portions of the *cis*-PT complex, yet focusing at the inter-subunit interface.

**Figure 2.**
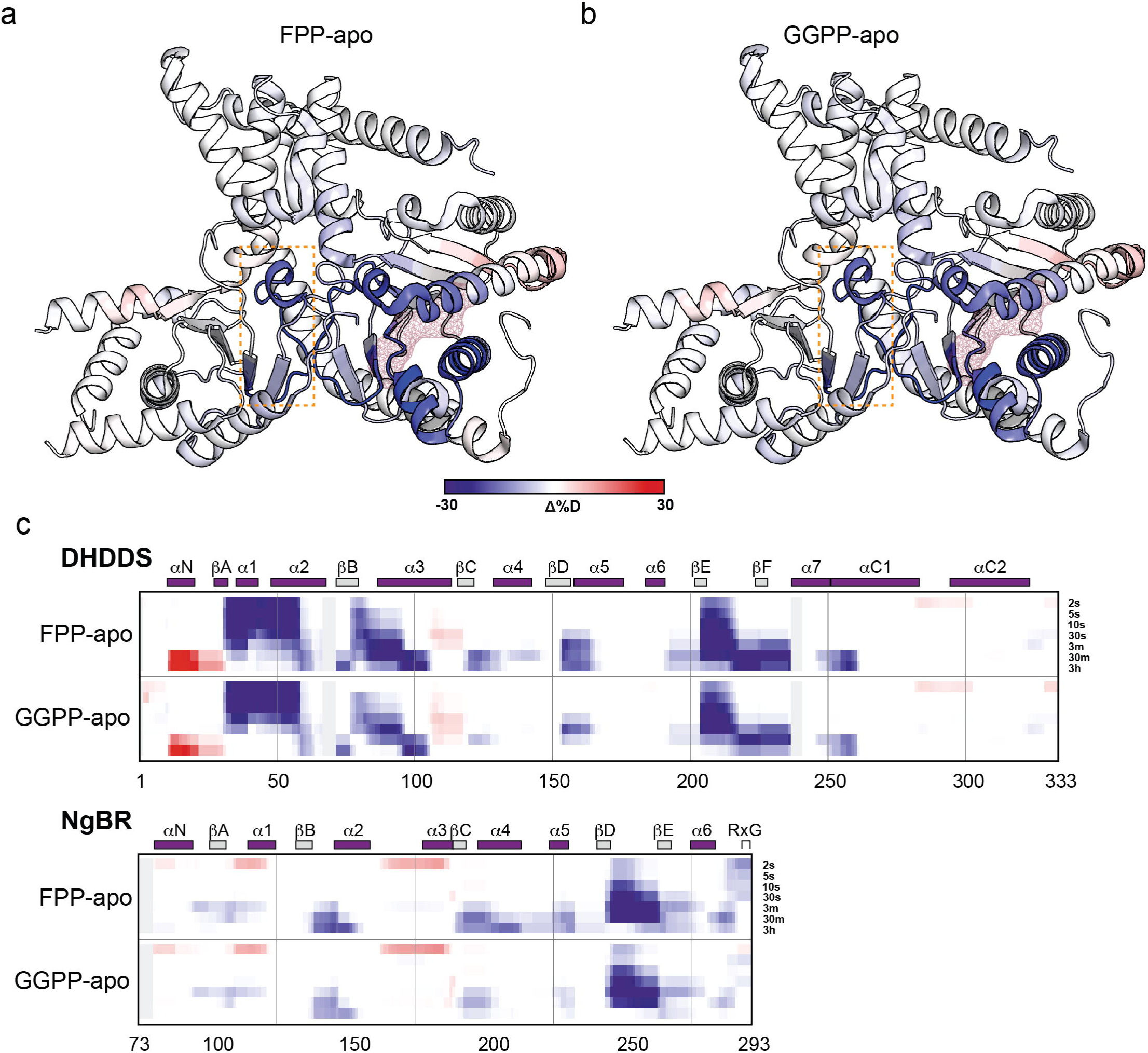
HDX-MS of h*cis*-PT reveals extensive dynamic response to active site occupancy state. (a,b) Cartoon representation of the differences in deuteration levels between FPP-(a) or GGPP-bound (b) and the apo states, following 30 seconds of incubation with D_2_O. Data are mapped onto the structure of FPP-bound h*cis*-PT (PDB 6Z1N). The red mesh indicates the position of the bound substrate. The dashed red rectangle frames the NgBR^βD-βE^ region. **(c)** Heatmap representations of the HDX differences between FPP- or GGPP-bound h*cis*-PT and the apo state at the indicated time points for DHDDS (upper panel) and NgBR (lower panel). Secondary structure elements are shown above the heat maps.

### Systematic functional mapping of NgBR interfacial region

The HDX-MS results suggest that the NgBR^βD-βE^ loop at the inter-subunit interface is a hub for allosteric communication (Fig. 3a)^36,37^. This is consistent with the high degree of evolutionary conservation at this region (Supplementary Fig. 1). To determine the functional contribution of specific residues within this region to h*cis*-PT enzymatic activity, we performed high-throughput alanine-scanning mutagenesis using a well-established yeast complementation platform (Fig. 3b)^22,27^. In this approach, the triple deletion KG405 strain, lacking the homologous yeast h*cis*-PT genes (τι*rer2*, τι*srt1*, τι*nus1*)^22^, is co-transformed with vectors containing DHDDS or NgBR and maintained under positive and negative selections. Only transformants complemented with functional h*cis*-PT subunits can thrive under these conditions^27^. Of the 19 positions examined, a single mutant, NgBR^S249A^, resulted in a lack of complementation, suggesting a functional perturbation prohibiting yeast growth.

**Figure 3.**
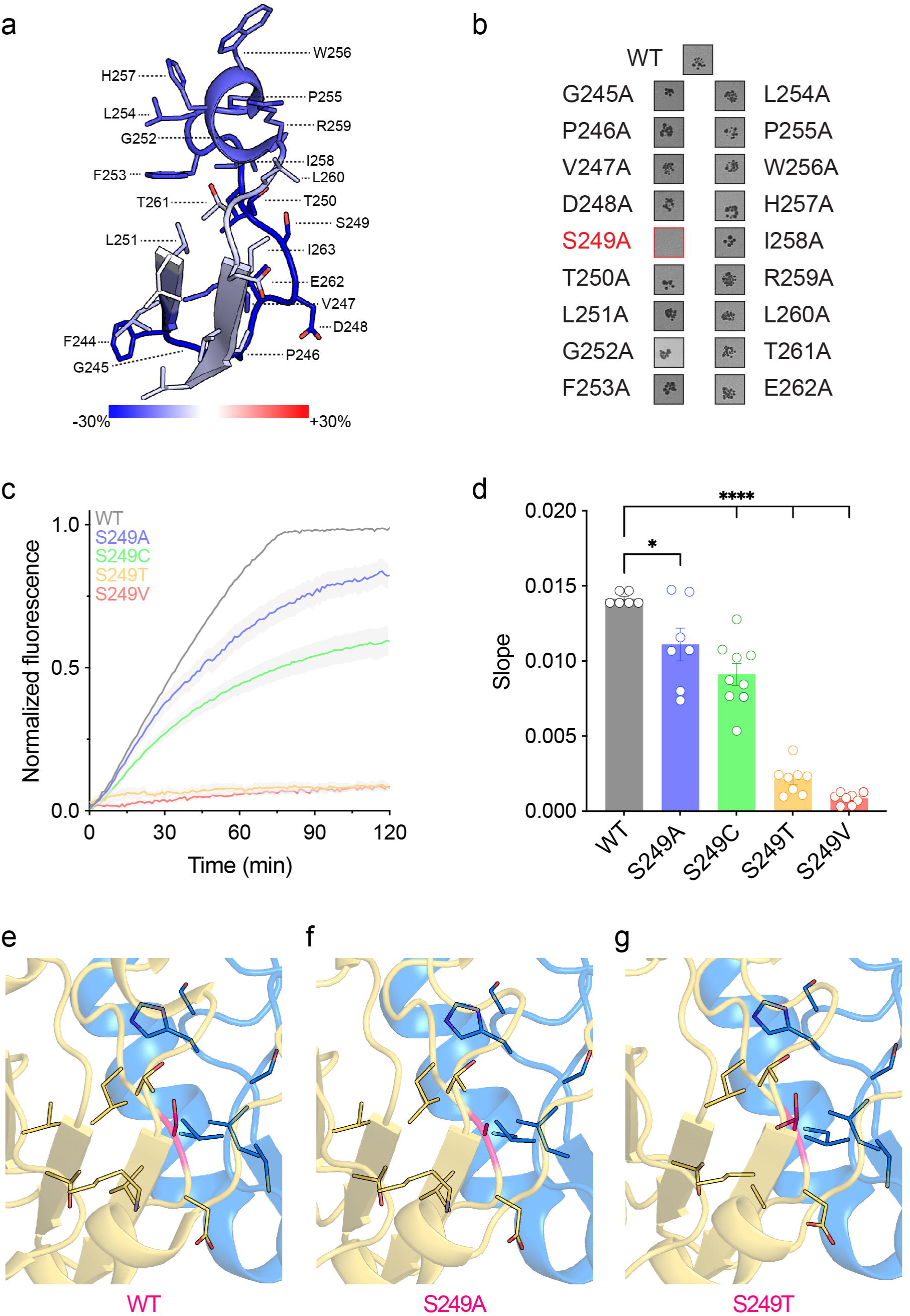
NgBR^S249^ is crucial for h*cis*-PT activity. **(a)** Zoomed perspective of the HDX difference between the FPP and apo forms at the NgBR^βD-βE^ region. **(b)** Yeast complementation-based high-throughput alanine scanning of positions 244-262 at the NgBR^βD-βE^ region.Only S249A failed to support yeast growth. **(c)** Average time-dependent increase in MANT-O-GPP fluorescence in the presence of h*cis*-PT harboring NgBR^WT^ versus the indicated S249 mutants. Data are presented as mean ± SEM (n = 6-9). **(d)** The initial slope of the fluorescence curves, reflecting the initial reaction rate (One-way ANOVA followed by Dunnett’s multiple comparisons test; * *P*<0.05, **** *P*<0.0001). **(e-g)** Inter-subunit interface zoomed view of hcis-PT containing NgBR^WT^ (e), NgBR^S249A^ (f), and NgBR^S249T^ (g) crystal structures. NgBR and DHDDS are colored in yellow and blue, respectively, and position 249 in pink.

To delineate the contribution of NgBR^S249^ to DHDDS activity modulation, we further mutated this position to residues with different physico-chemical properties (A, T, V, C) to probe the significance of both residue volume and polarity on NgBR regulation of h*cis*-PT function. Following purification, the mutants were subjected to *in vitro* fluorescence-based enzyme kinetics analyses. This assay utilizes (2E,6E)-8-O-(N-methyl-2-aminobenzoyl)-3,7-dimethyl-2,6-octandien-1-pyrophosphate (MANT-O-GPP)^38^, a fluorescent FPP analog. MANT-O-GPP exhibits an increase in fluorescence upon chain elongation by h*cis*-PT, allowing the monitoring of catalytic activity via fluorescence spectroscopy (Fig. 3c)^27,38^. Importantly, using this approach, we observed a dramatic reduction in catalytic activity upon all NgBR^S249^ substitutions tested. Specifically, while NgBR^S249A^ and NgBR^S249C^ exhibited a moderate decrease in catalytic activity, judged from the amplitude and slope of the fluorescence increase, the NgBR^S249V^ and NgBR^S249T^ mutants nearly eliminated the activity (Fig. 3c, d). Finally, consistent with the narrow conformational space sampled by the crystal structures obtained to date, crystallographic analysis of substrate-bound NgBR^S249A^ and NgBR^S249T^ did not reveal any local or global conformational alterations (root mean square deviation (RMSD) = 0.15 Å and 0.33 Å, respectively, compared to PDB 7PAX) (Fig. 3e-g). These results indicate that the side-chain volume rather than the polarity at position NgBR^249^ plays a crucial role in conveying allosteric information necessary for DHDDS activity regulation by NgBR, without provoking global conformational changes.

### hcis-PT allosteric network analysis reveals inter-subunit communication paths

The HDX-MS analysis pinpointed the NgBR^βD-βE^ loop as a functional hub, with the yeast complementation and enzymatic activity assays strongly supporting a unique role for the conserved NgBR^S249^ in the modulation of DHDDS activity. As the high-resolution structures of the mutant h*cis*-PT showed limited conformational alterations, we posited that transient dynamic coupling between NgBR^S249^ and catalytic DHDDS residues might be at play. To explore this possibility, we performed triplicates of 250 nsec all-atom MD simulations of the WT and NgBR^S249V^ complexes (Supplementary Fig. 2). All the simulations performed here have reached convergence, as reflected by the plateau in RMSD values (Supplementary Fig. 2a,c). In addition, the per-residue root mean square fluctuation (RMSF) values, representing its spatial fluctuation relative to its average position along the simulation, are largely unaffected by the mutation (Supplementary Fig. 2b,d). Next, the simulation trajectories were subjected to network analysis (Fig. 4)^39^. This technique was also successfully applied in the study of activation^40^ and inactivation ^41^ of ion channels, producing results in agreement with patch clamp experiments^42^ and suggesting possible applications for drug design^43^. In this approach, the protein is represented as a network, where nodes and arcs represent the residues and their interactions, respectively (see Methods). Each arc is assigned a weight that quantifies the efficiency of motion propagation between the two residues flanking the arc. To delineate the allosteric paths, we defined the NgBR^S249^ region as the source, while the sink region is represented by one of the residues of the catalytic pocket and its nearest neighbors. Next, we computed the paths of minimal length on the graph between all the residues of the source region and all the residues of the sink region^44^, where the path length is the sum of the weights of the arcs along the path. Finally, a Centrality Index (CI), representing the fraction of minimal paths the residue participates in, is computed for each residue. Residues with a high CI value are at the crossroads of many pathways and are thus likely to act as hubs in the communication network.

**Figure 4.**
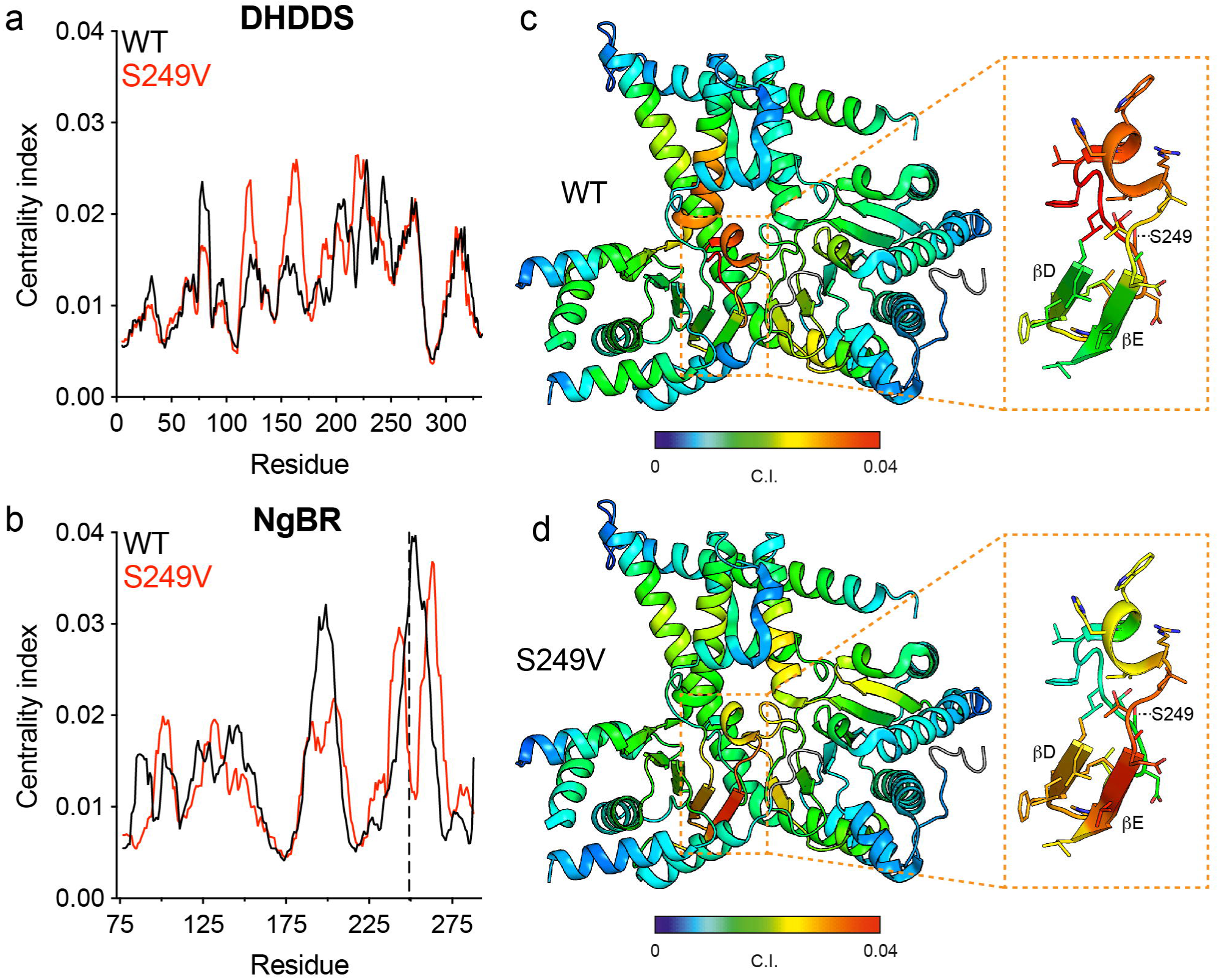
Allosteric path analysis highlights the NgBR^βD-βE^ region as a hub inter-subunit communication. (a,b) Per-residue global CI analysis, taking into account all possible paths connecting every residue to all other residues, of DHDDS (a) and NgBR (b), obtained from MD simulations of h*cis*-PT harboring NgBR^WT^ (black) or NgBRS^249V^ (red). **(c,d)** Global CI heatmaps for h*cis*-PT harboring NgBR^WT^ (c) or NgBR^S249V^ (d) projected on the structure in cartoon representation. The provided blow-ups of the NgBR^βD-βE^ region highlight the interference induced by the NgBR^S249V^ mutation.

Strikingly, the global CI, computed on all possible paths connecting every residue to all other residues, exhibits its highest peak in the NgBR^βD-βE^ loop (Fig. 4a,b). This may explain the dramatic impact of NgBR^S249^ mutant because this position forms a critical crossroads where the largest number of routes pass. Conversely, in the NgBR^S249V^ mutant, this region finds itself at the bottom of the valley between the peak of strand βD and the shifted peak of the NgBR^βD-βE^ loop (that also incorporates the βE region) (Fig. 4c,d). Thus, the NgBR^S249V^ mutation dramatically abates the CI, that is, much fewer pathways transit through this position.

Based on this analysis, we identified allosteric paths originating from NgBR^S249^ and ending at various catalytic positions within the DHDDS active site (Fig. 5a). Then, we selected six positions in DHDDS, spanning the βE and βF strands, as well as the βF-α7 linker, for validating the functional significance of the identified paths. All mutants affected enzymatic activity in line with our network analysis (Fig. 5b,c). Interestingly, most mutants resulted in decreased activity, as reflected by reduced fluorescence amplitude and/or slope, one mutant, DHDDS^F227A^, demonstrated enhanced activity. Together, our results expose allosteric communication pathways between the inter-subunit interface and the active site of DHDDS.

**Figure 5.**
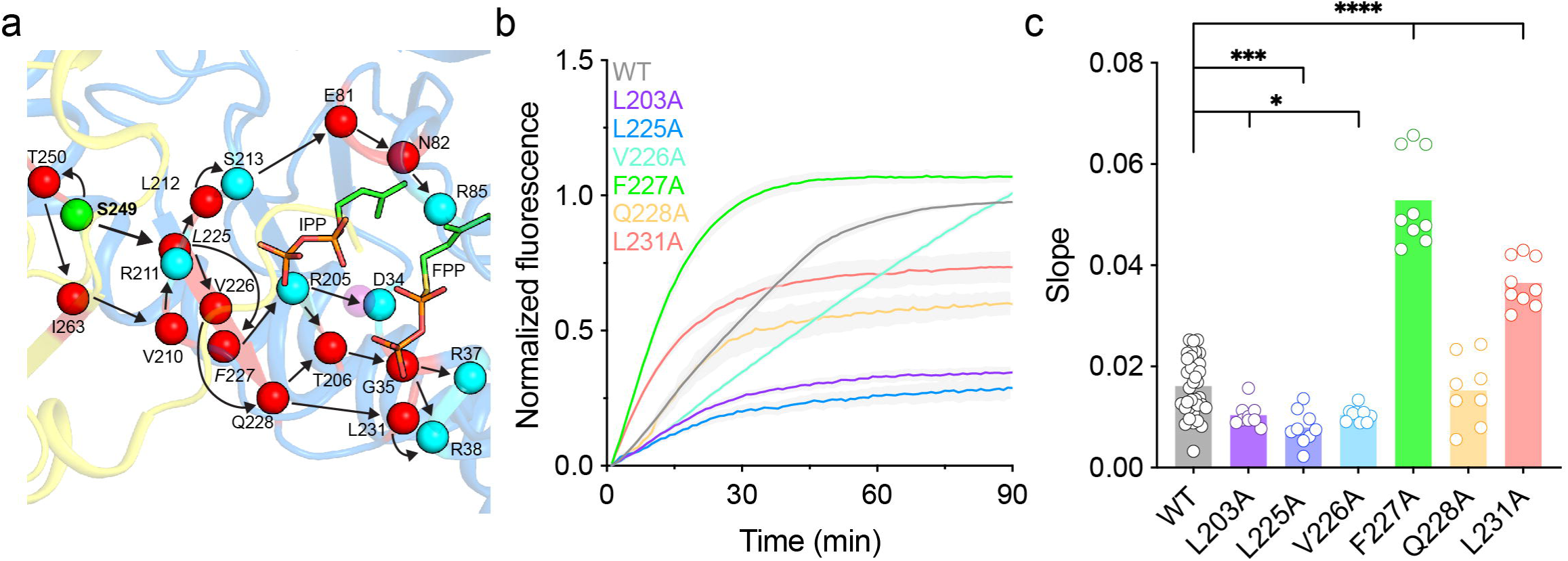
Inter-subunit allosteric communication paths perturbations alter h*cis*-PT function. **(a)** Network analysis of allosteric communication path trajectories. Originating from NgBR^S249^ (green sphere), the paths connecting the predicted network (red spheres) and catalytic (cyan spheres) residues are highlighted (arrows). Note that in some cases, network and catalytic residues overlap. **(b)** Average time-dependent increase in MANT-O-GPP fluorescence in the presence of h*cis*-PT harboring DHDDS^WT^ versus the indicated allosteric path mutants. Data are presented as mean ± SEM (n = 6-9). (c) The initial slope of the fluorescence curves, reflecting the initial reaction rate (One-way ANOVA followed by Dunnett’s multiple comparisons test; * *P*<0.05, *** *P*<0.001, **** *P*<0.0001).

## Discussion

h*cis*-PT is a membrane-associated enzymatic complex that produces long-chain isoprenoids crucial for numerous cellular processes. Importantly, the long-chain subclass of prenyltransferases consists of heteromeric complexes composed of soluble-catalytic and membrane-anchored auxiliary subunits^1^. Given the hydrophobicity of their polyprenol products, it was suggested that membrane association may contribute to ER membrane dolichol enrichment for downstream N-linked glycosylation. Indeed, close vicinity to the membrane is expected to enhance catalytic efficiency and proper product placement. However, NgBR was also shown to dramatically enhance the catalytic activity of DHDDS^26,28,45^, even without its transmembrane region, suggesting a regulatory role beyond mere membrane anchoring^27^. Here, by employing a combined experimental-computational approach, we unravel the molecular mechanisms underlying the allosteric regulation of DHDDS activity by NgBR.

Previously, to obtain high-resolution insights into the catalytic mechanism of h*cis*-PT, we obtained high-resolution structures of the complex in discrete states along its catalytic cycle^27^, revealing no significant conformational changes at NgBR in different substrate/product-bound states. To expose possible reciprocal conformational changes in NgBR related to DHDDS active-site occupancy, we crystallized h*cis*-PT in its apo-form. This structure revealed only minor conformational changes focused exclusively on the active site (Fig. 1). Only local rearrangements in key active site residues and the distal C-terminal tail of NgBR were observed. However, the crystalline environment, as well as the cryogenic temperature used in the diffraction experiment, result in conformational convergence and prohibit a complete sampling of the conformational space^46^.

Given the lack of major structural rearrangements, we asked whether changes in the protein motions mediate the striking effect of NgBR on the enzymatic turnover rate^47^. Along this line, previous studies have repeatedly shown that allosteric control of enzymatic function could elude detection due to the lack of associated large-scale conformational transitions^48^. HDX-MS is increasingly used to expose long-range dynamic perturbations induced by the binding of cofactors, substrates, or inhibitors to enzymes^49–52^, successfully probing the dynamic realm of protein function, which is often unattainable by high-resolution static structural approaches^53^. Therefore, we subjected h*cis*-PT to HDX-MS analysis in the apo and substrate-bound forms, revealing widespread dynamic changes in solution, spanning throughout the complex and propagating well beyond the active site (Fig. 2). Specifically, this analysis highlighted the interface between NgBR and DHDDS as a potential crossroad for functional inter-subunit communication. In support of the possible functional role of the inter-subunit interface, a comparison of the heteromeric interface with the modeled homodimeric DHDDS structure revealed a higher number of interacting residues, leading to a more intimate association between the subunits.

We identified NgBR^S249^ as a critical node for relaying the allosteric signal between the subunits within the inter-subunit interface. Indeed, this conserved position did not tolerate the introduction of mutations, resulting in markedly reduced catalytic activity without causing major structural perturbations (Fig. 3), further underscoring the dynamic nature of inter-subunit communications in the h*cis*-PT complex. Comparing different substitutions (A, C, T, V), the significance of side-chain volume for proper enzymatic function emerged. Consistent with this observation, complexes harboring NgBR^S249T^ exhibited relatively poor diffraction (Table 1), while those harboring NgBR^S249V^ were calcitrant to crystallization altogether, possibly resulting from interface destabilization. Importantly, the NgBR^S249T^ and NgBR^S249V^ mutations result in activity reminiscent of the homodimeric DHDDS complex^45^, surpassing the deleterious effect of the disease-causing NgBR^R290H^, suggesting that the NgBR^βD-βE^ loop at the inter-subunit interface plays a crucial role in inter-subunit communication.

**Table 1.**
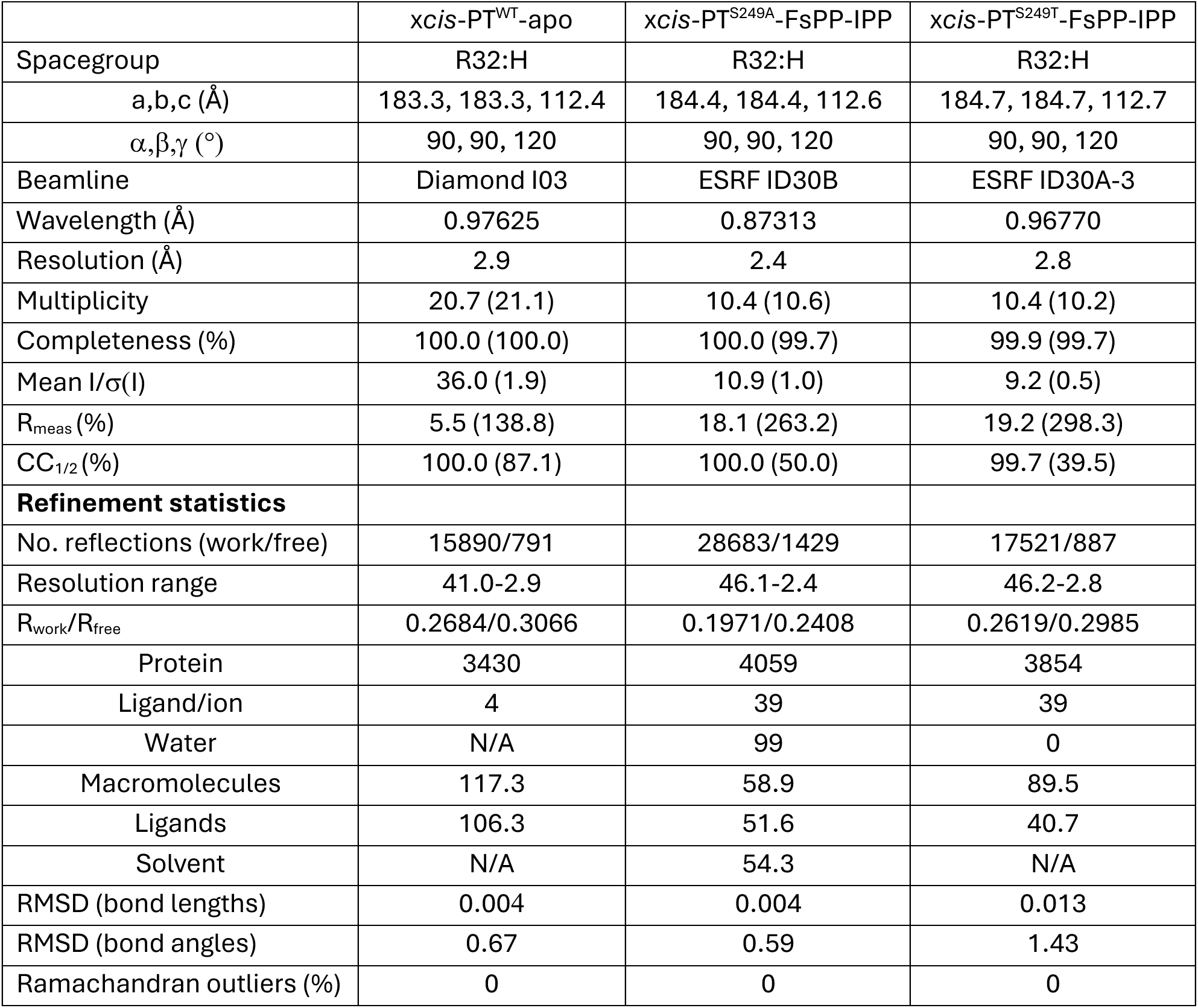
Crystallographic statistics.

|  | *xcis*-PT<sup>WT</sup>-apo | *xcis*-PT<sup>S249A</sup>-FsPP-IPP | *xcis*-PT<sup>S249T</sup>-FsPP-IPP |
| --- | --- | --- | --- |
| Spacegroup | R32:H | R32:H | R32:H |
| a,b,c (Å) | 183.3, 183.3, 112.4 | 184.4, 184.4, 112.6 | 184.7, 184.7, 112.7 |
| $\alpha, \beta, \gamma$ (°) | 90, 90, 120 | 90, 90, 120 | 90, 90, 120 |
| Beamline | Diamond I03 | ESRF ID30B | ESRF ID30A-3 |
| Wavelength (Å) | 0.97625 | 0.87313 | 0.96770 |
| Resolution (Å) | 2.9 | 2.4 | 2.8 |
| Multiplicity | 20.7 (21.1) | 10.4 (10.6) | 10.4 (10.2) |
| Completeness (%) | 100.0 (100.0) | 100.0 (99.7) | 99.9 (99.7) |
| Mean I/$\sigma$(I) | 36.0 (1.9) | 10.9 (1.0) | 9.2 (0.5) |
| R<sub>meas</sub> (%) | 5.5 (138.8) | 18.1 (263.2) | 19.2 (298.3) |
| CC<sub>1/2</sub> (%) | 100.0 (87.1) | 100.0 (50.0) | 99.7 (39.5) |
| **Refinement statistics** |  |  |  |
| No. reflections (work/free) | 15890/791 | 28683/1429 | 17521/887 |
| Resolution range | 41.0-2.9 | 46.1-2.4 | 46.2-2.8 |
| R<sub>work</sub>/R<sub>free</sub> | 0.2684/0.3066 | 0.1971/0.2408 | 0.2619/0.2985 |
| Protein | 3430 | 4059 | 3854 |
| Ligand/ion | 4 | 39 | 39 |
| Water | N/A | 99 | 0 |
| Macromolecules | 117.3 | 58.9 | 89.5 |
| Ligands | 106.3 | 51.6 | 40.7 |
| Solvent | N/A | 54.3 | N/A |
| RMSD (bond lengths) | 0.004 | 0.004 | 0.013 |
| RMSD (bond angles) | 0.67 | 0.59 | 1.43 |
| Ramachandran outliers (%) | 0 | 0 | 0 |

While studies of the h*cis*-PT complex offer a wealth of molecular information, it remains challenging to integrate the static crystallographic structures with the dynamic information obtained by HDX-MS, which lacks spatial resolution. We addressed this gap by performing allosteric communication path analyses of MD simulation trajectories. These analyses suggest that the h*cis*-PT complex constitutes a specialized fold primed at efficiently conducting cross-subunit allosteric information (Fig. 4). Specifically, the global CI exhibited its highest peak at the NgBR^βD-βE^ loop, in agreement with the marked dynamic response of this region to substrate binding at the DHDDS active site. Indeed, mutational analysis of select residues along the paths interconnecting NgBR^S249^ and DHDDS catalytic residues confirmed their functional significance (Fig. 5).

Altogether, our work exposed hidden links to active site residues, providing mechanistic insights into NgBR-mediated functional regulation of DHDDS. We show that intimate inter-subunit structural and functional interactions are crucial for proper h*cis*-PT activity. This functional modulation stems from protein dynamics encoded within the protein fold. Importantly, as h*cis*-PT is a central metabolic node in every cell, playing pivotal roles in numerous cellular processes and is crucial for human health, our results generate new opportunities for guided design of allosteric activators for the treatment of h*cis*-PT-related diseases.

## Methods

### Protein expression and purification

*E. coli* T7 express competent cells were co-transformed with DHDDS (residues 1-333) and NgBR (residues 73-293) (h*cis*-PT). For crystallization trials, NgBRτι167-175 was used (x*cis*-PT) as previously described^26,27^. Cells were grown in Terrific Broth medium at 37 °C until reaching OD_600nm_ = 0.6 and induced at 16 °C by adding 0.5 mM isopropyl β-D-1-thiogalactopyranoside (IPTG). Proteins were expressed at 16 °C for 16-20 hr, harvested by centrifugation (∼5,700 xg for 15 min), and then resuspended in a buffer containing 20 mM 4-(2-hydroxyethyl)-1-piperazineethanesulfonic acid (HEPES), pH 7.5, 150 mM NaCl, and 1 mM tris(2-carboxyethyl)phosphine (TCEP) and 0.02% (w/v) triton X-100, supplemented with 1 μg/ml DNase I and a protease inhibitor mixture. Resuspended cells were homogenized and disrupted in a microfluidizer. Soluble proteins were recovered by centrifugation at ∼ 40,000×g for 45 min at 4 °C. Overexpressed proteins were purified on a HisTrap HP column, followed by purification on a Strep-Tactin column and TEV protease cleavage of the purification tags and TRX fusions. The reaction mixture was concentrated and loaded onto a Superdex-200 preparative size-exclusion column pre-equilibrated with 20 mM 4-(2-hydroxyethyl)-1-piperazineethanesulfonic acid (HEPES), pH 7.5, 150 mM NaCl, 1 mM tris(2-carboxyethyl)phosphine (TCEP). Purified proteins were flash-frozen in liquid nitrogen and stored at -80 ° C until use. Protein purity was >95%, as judged by SDS-PAGE.

### Crystallization and structure determination

All crystallization trials were performed at 19 °C using the sitting drop vapor diffusion method. Crystals of apo x*cis*-PT were obtained by mixing protein solutions at ∼16 mg/mL with 0.1 M LiCl, 0.05 M Glycine pH 9.5, and 40% (v/v) PEG 400. Crystals were cryoprotected and data were collected at 100 K at the Diamond Light Source (DLS; Oxfordshire, United Kingdom). The x*cis*-PT mutant complexes were crystallized by mixing protein solutions at ∼7.5 mg/mL in the presence of 0.5 mM MgCl_2_, 0.75 mM FsPP, and 1.5 mM IPP with 0.1 M Tris pH 8.5, 0.1 M KCl, 39% (v/v) PEG 400, (DHDDS^L225A^ and DHDDS^L225N^), 0.1 M Tris pH 9, 1 M Glycine, 0.1M KCl, 41% (v/v) PEG 400 (NgBR^S249A^) or 0.1 M Hepes pH 8, 30% Pentaerythritol ethoxylate, 4% PVP K15, 30% (w/v) 1,6-Hexanediol (NgBR^S249T^). Crystals were cryoprotected and data were collected at 100 K at the European Synchrotron Radiation Facility (ESRF; Grenoble, France). Integration, scaling and merging of the diffraction data were done with the XDS program^54^. All the structures were solved by automated molecular replacement using Phaser^55^ with the structure of h*cis*-PT in complex with FsPP and IPP (PDB 7PAX) as a search model. Iterative model building and refinement were carried out in PHENIX ^56^ with manual adjustments using COOT^57^. Ramachandran analysis was performed using MolProbity ^58^. Data collection and refinement statistics are presented in Table 1. Structural illustrations were prepared with PyMOL 3.0 (Schrödinger LLC).

### Yeast complementation

The *S. cerevisiae* strain K405, lacking the yeast DHDDS and NgBR homologs (Δ *nus1*, Δ *rer2*, Δ *srt1*) and carrying the Gl*cis*-PT on a plasmid bearing the *URA3* selection marker was used ^22^. To analyze NgBR interdomain interface mutants, cells were transformed using the standard lithium-acetate method with the DHDDS.GW1 and NgBR.GW2 (WT or mutants thereof) plasmids, bearing leucine and methionine selection markers, respectively. Empty vectors were used as negative controls, and Gl-*cis*PT.GW1 was used as a positive control. Following transformation, cells were plated on solid synthetic defined medium lacking uracil, methionine, and leucine and incubated for 48-72 hrs at 30 °C. Next, single colonies were inoculated into liquid synthetic defined medium lacking uracil, methionine, and leucine, incubated overnight at 30 °C using a shaker, and diluted to reach OD_600_ = 1. For phenotypic analysis, transformants were spotted on solid synthetic defined complete medium supplemented with 0.1% (w/v) FOA plates, and incubated at 30°C for 5 days. Synthetic defined medium was made of 0.67% (w/v) yeast nitrogen base and 2% (w/v) supplemented with auxotrophic requirements. For solid medium, agar was added at a 2% (w/v) final concentration.

### Enzyme kinetics

MANT-O-GPP was synthesized as previously described ^38^. All fluorescence experiments were performed in triplicate using an RF-8500 spectrofluorometer (Jasco, Japan). Briefly, purified h*cis*-PT (0.2 μM) and MANT-O-GPP (5 μM) were premixed in fluorescence buffer, consisting of 20 mM Tris-HCl (pH 7.5), 150 mM NaCl, 10 mM β-mercaptoethanol, and 0.5 mM MgCl_2_. To initiate the reaction, IPP (100 μM) was quickly added and mixed, and fluorescence time course measurements (n ≥ 6 replicates) were conducted immediately using Ex = 352 nm and Em = 425 nm, excitation and emission wavelengths, respectively. Data were normalized and plotted using Prism GraphPad 9.0.1.

### HDX-MS

Hydrogen-deuterium exchange was followed for hcis-PT in the apo, FPP, or GGPP-bound states. Proteins in 20 mM HEPES pD 7.5, 150 mM NaCl, 1mM TCEP were supplemented with 0.5 mM EDTA in the apo conditions or with 5 mM MgCl_2_ and 6-molar excess of FPP or GGPP for the ligand-bound states. The exchange was done manually at 4 °C. It was initiated by a 10-fold dilution of 20 μM protein solution (for FPP/GGPP the ligands were 120 uM) into a D_2_O-based buffer (pD 7.5) and aliquots were collected after 2 s*, 5 s, 10 s*, 30 s, 3 m*, 30 m, 3 h* (time point denoted with * were replicated). The exchange was quenched with 0.5 M glycine-HCl, pH 2.5 in a 1:1 ratio, samples were frozen in liquid nitrogen and stored at -80 °C. Fully deuterated samples were analyzed to correct the data for back-exchange during the analysis. Before the analysis, each sample was thawed and injected onto an LC system. Proteins were digested online (200 µL/min, 0.4% formic acid in water) on an immobilized pepsin/nepenthesin-2 column (bed volume 66 µL), and the peptides were desalted on a trap column (SecurityGuard ULTRA Cartridge UHPLC Fully Porous Polar C18, 2.1 mm ID; Phenomenex). Digestion and desalting took 3 minutes. Next, the peptides were separated on an analytical column (Luna Omega Polar C18, 1.6 µm, 100 Å, 1.0×100 mm; Phenomenex) by water-acetonitrile gradient (5%–45% B within 7 min; solvent A: 0.1% FA in water, solvent B: 0.1% FA/2% water in ACN). The solvents were delivered by 1290 Infinity II LC System (Agilent Technologies, Waldbronn, Germany) at 40 µL·min^-1^. The whole setup was cooled to 0 °C to minimize deuterium loss. The LC system was interfaced with an ESI source of 15 FT-ICR MS (solariX XR, Bruker Daltonics, Bremen, Germany) operating at MS1 mode with 1M data-point acquisition. The LC-MS data were peak-picked in Data Analysis (Bruker Daltonics) and further processed by the DeutEx software^59^. Data visualizations were done by MSTools (http://peterslab.org/MSTools/index.php)^60^ and PyMol. Correction for back-exchange was done as described previously^61^.

For peptide identifications, the same LC setup as described above was used, but the LC was connected to a timsToF Pro (Bruker Daltonics, Bremen, Germany). LC-MS/MS analysis employing PASEF-based data-dependent (CID) analysis was done on non-deuterated samples. The data were exported to *.mgf file and searched against a custom-built database using MASCOT (v. 2.7, Matrix Science, London, United Kingdom). The database combined cRAP.fasta (https://www.thegpm.org/crap/) with the sequences of NgBR and DHDDS constructs and the proteases used. Search parameters were as follows – no modifications included, precursor tolerance 10 ppm, fragment ion tolerance 0.05 Da. Decoy search was enabled and FDR set to <1% and IonScore >20. All mass spectrometry data have been deposited to the ProteomeXchange Consortium via the PRIDE partner repository with the dataset identifier PXD063722^62^.

### MD simulations

All the simulations were performed using the Schrödinger Maestro release 2022-3 (Schrödinger LLC). First, missing loops were added to the structure of h*cis*-PT in complex with FPP (PDB 6Z1N) using MODELLER^63^, and the NgBR^S249V^ mutation was introduced using Schrödinger Maestro release 2022-3 (Schrödinger, LLC). Next, the ligands were removed and the structure was prepared using the Protein Preparation Wizard. Missing hydrogen atoms were added considering a pH value of 7.2 ± 1.0, followed by optimization of the hydrogen bond network. The system setup tool was used to solvate the systems using the TIP3P solvent model. Potassium or chloride ions were added to neutralize the charge and to obtain a final salt concentration of 150 mM. All MD simulations were performed using Desmond with the OPLS4 force field ^64^. The simulations were conducted under a Langevin temperature and pressure control, using periodic boundary conditions with particle-mesh Ewald (PME) electrostatics with a 9 Å cutoff for long-range interactions. The systems were equilibrated using the default relaxation protocol. Finally, the production simulations were carried out for 250 ns with a constant pressure of 1 atm and a constant temperature of 300 K starting from a random seed. The results were manually inspected using the Maestro suite.

### Allosteric path analysis

In network analysis, the protein is modeled as a network where nodes and arcs represent residues and interaction between residues, respectively. Each residue is assigned a weight that accounts for both connectivity and correlation between the two residues. In particular, the weights are defined as: *G_ij_*= -log(*C_ij_*\**I_ij_*) where *C_ij_* is a semi-binary contact map and *I_ij_* is mutual information^65^. With this choice small values of our metric correspond to pairs of residues in contact and with a highly correlated motion. By contrast, high values of this ”information distance” represent residues whose motion is not correlated and/or not in contact with one another. In other words the weight can be considered as a proxy for causation. We do not know whether residue *i* drives the dynamics of residue *j* (or *vice versa*), but if the two residues are in contact and they move in a coordinated way (small value of the weight), it is likely that a causal link between them exists. The pathways identified with this approach can thus be considered as kinematic chains, like the tiles of the domino game.

Contacts are normally described through a binary contact map where a value 1 labels a pair of residues in contact while a value 0 indicates that there is no contact. Conversely a semi-binary contact map assigns a value 1 to a pair of residues below a distance cutoff *c*=7.0 Å and a gaussian decreasing value exp(-(*d*^2^-*c*^2^)/2*σ*^2^) if the distance *d* exceeds the threshold. This gaussian kernel is then averaged over all the frames of the simulation. The semi-binary contact maps allow to account for those pairs of residues whose distance is just above the cutoff.

The correlation in the motion of a pair of residues *i*, *j*, is quantified through Mutual Information. In Information Theory the Mutual Information quantifies the decrease in the uncertainty of a random variable *X* due to the knowledge of another random variable *Y* and thus quantifies the coupling between the two variables. In our case, the two variables represent the displacements *d_i_*, *d_j_* of two side chains from their average position:

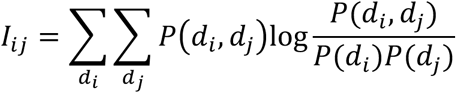

Mutual information thus quantifies the correlation between the motion of two residues. It is notable that while the customarily used correlation coefficient can only capture linear correlations, the mutual information can also detect higher-order correlations.

Once the network representation is built, we need to study how motion propagates between different districts of the protein. For instance, let us consider the communication between the βD-βE-loop of NgBR subunit and the βE-βF-loop of the catalytic subunit (Supplementary Fig. 3). In order to do so, we choose a central residue in both the source and sink regions. We then build a sphere of radius 7.0 Å centered on the two key residues, and we identify the residues located inside the two spheres in at least 75% of the frames of our trajectory. Using Dijkstra’s algorithm ^44^, we then compute the minimal paths between all pairs of residues belonging to the first and second regions. Finally, for each residue, we compute a CI, i.e., the fraction of minimal paths the residue sits on.

It is important to note that while the CI computed in the analysis of the pathways starting from NgBR^S249^ is local because it was computed only considering the pathways that connect the residues in the neighborhood of NgBR^S249^ with the residues in the neighborhood of the target, the CI plotted in Fig. 4a,b is global because it was computed considering all possible pathways connecting all residues to all other residues.

### Data Availability

Atomic coordinates and structure factors for the structures of x*cis*-PT^WT^-apo (PDB 9R0K), x*cis*-PT^S249A^-FsPP-IPP (PDB 9R08), and x*cis*-PT^S249T^-FsPP-IPP (PDB 9R0E), have been deposited in the Protein Data Bank. The mass spectrometry proteomics data have been deposited in the ProteomeXchange Consortium via the PRIDE^62^ partner repository with the dataset identifier PXD063722.

## Acknowledgments

We thank the staff of ID30A-3 and ID30B at the European Synchrotron Radiation Facility (ESRF), and I03 at the Diamond Light Source for assistance with diffraction experiments. This work was supported by the Israel Science Foundation (grant 1653/21 (Y.H.)), the Israel Cancer Research Fund grants 1289067 and (Y.H. and M.G.), the Israel Cancer Association grants 20230029 (Y.H. and M.G.), the Bi-National Science Foundation grant 2023190 (Y.H.), and the Kahn Foundation’s Orion project, Tel Aviv Sourasky Medical Center, Israel (M.G.). Support also came from the Claire and Amedee Maratier Institute for the Study of Blindness and Visual Disorders, Faculty of Medicine, Tel-Aviv University (Y.H. and M.G.). P.M. acknowledges support from MEYS/EU project OP JAK—Photomachines (CZ.02.01.01/00/22_008/0004624). Access to MS instrumentation was enabled by CIISB LM2023042 and ERDF “UP CIISB” (CZ.02.1.01/0.0/0.0/18_046/0015974) and the EU Horizon 2020 grant EU_FT–ICR_MS (731077). Support also came from National Recovery and Resilience Plan (NRRP), Mission 4, Component 2, Investment 1.1, Call for tender No. 1409 published on 14.9.2022 by the Italian Ministry of University and Research (MUR), funded by the European Union – NextGenerationEU – Project Title “The virtual EV (v-EV): A digital twin of extracellular vesicles for health and food’’ – CUP B53D23027530001 -Grant Assignment Decree No. 1389 adopted on 01/09/2023 by the Italian Ministry of Ministry of University and Research (MUR) (C.G. and A.G.).

## Author contributions

Conceptualization, M.G. and Y.H.; Methodology, C.G., A.G., P.M., M.G. and Y.H.; Investigation, S.K., C.G., L.A., P.V., A.G., P.M., M.G., and Y.H.; Formal Analysis, C.G., P.V., P.M., M.G., and Y.H.; Writing – Original Draft, M.G. and Y.H.; Writing – Review and Editing, S.K., C.G., L.A., P.V., A.G., P.M., M.G., and Y.H.; Supervision, M.G. and Y.H.; Funding Acquisition, A.G., P.M., M.G., and Y.H..

