## Supplementary information for "Structural mechanisms of allosteric regulation in the human *cis*-prenyltransferase complex"

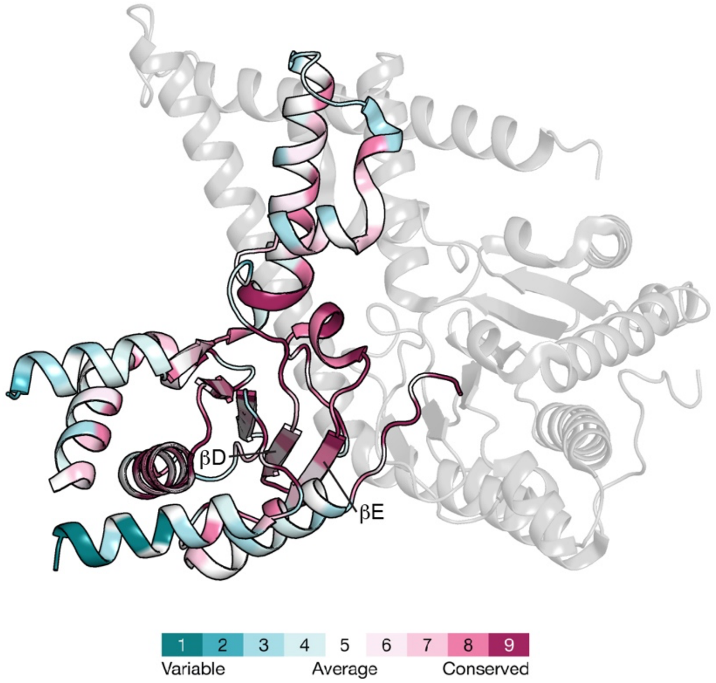


**Supplementary Figure 1. Conservation analysis of NgBR.** Conservation scores were calculated using the ConSurf server (https://consurf.tau.ac.il/consurf_index.php) and are projected on the NgBR subunit.

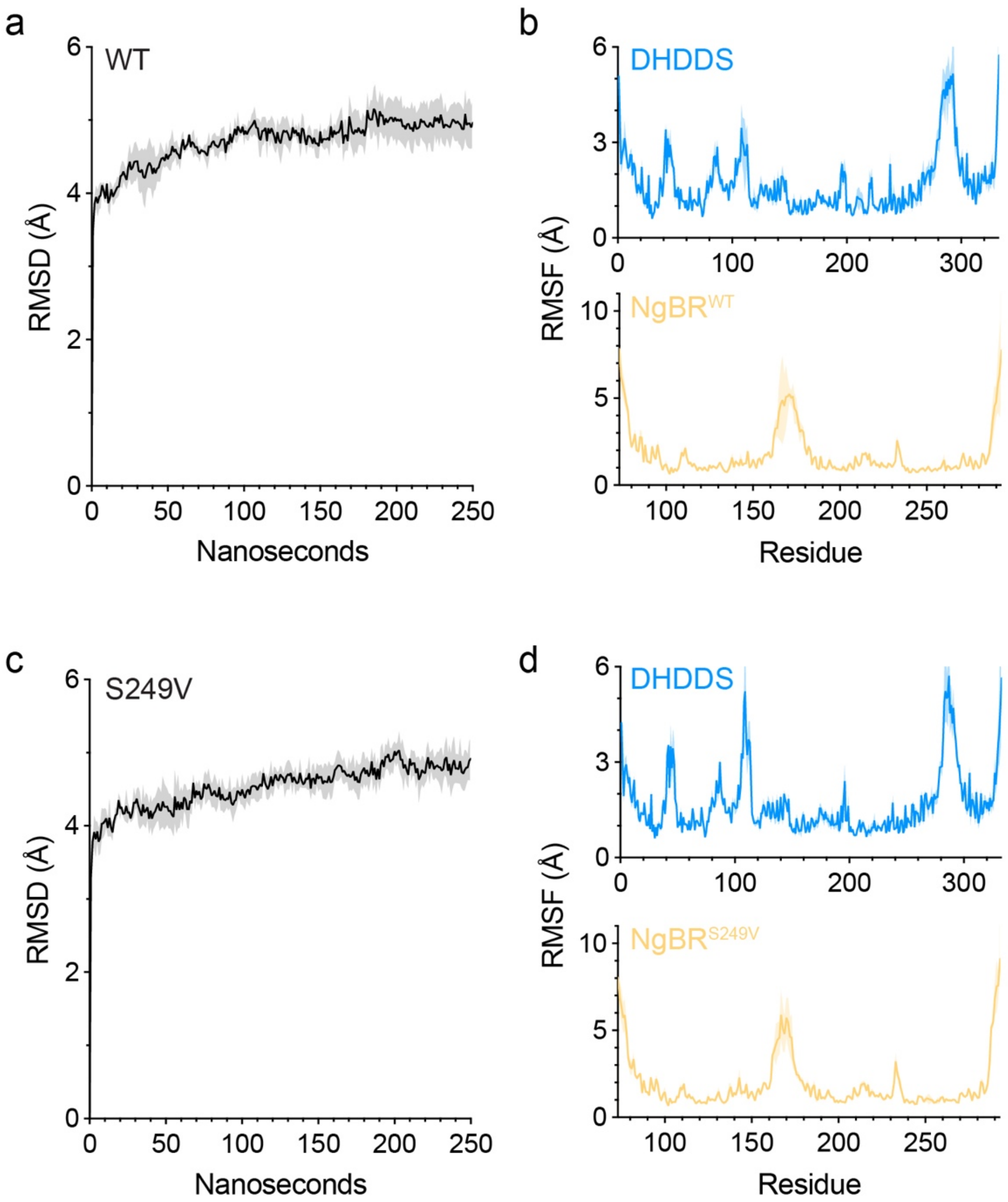


**Supplementary Figure 2. Molecular dynamics simulations of h*cis*-PT. (a,c)** Average root mean square deviation (RMSD) of h*cis*-PT containing $NgBR^{WT}$ (a) or $NgBR^{S249V}$ (c) along the simulation trajectories performed (n=3). The plateau indicates that the simulations reached convergence. The shaded area indicates the standard deviation. **(b,d)** Average root mean square fluctuation (RMSF), representing the per-residue spatial fluctuation relative to their average position along the simulation trajectories, of h*cis*-PT containing $NgBR^{WT}$ or $NgBR^{S249V}$ (d).

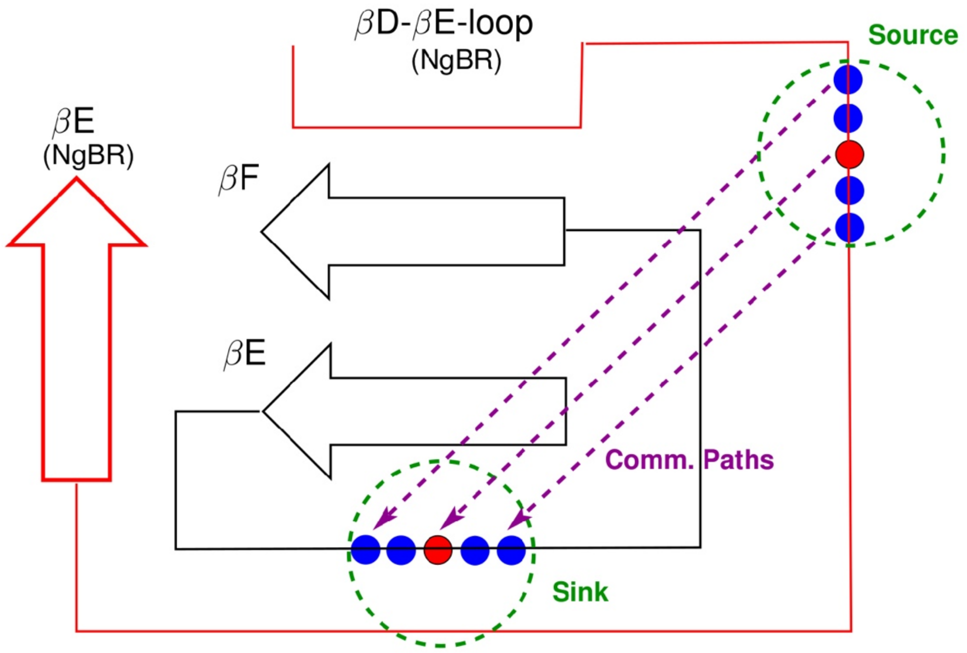


**Supplementary Figure 3. Cartoon representation of the pathway analysis method.** Two fragments of the DHDDS and NgBR subunits are shown in black and red, respectively. The source and sink regions are spheres of radius 7.0 Å centered on the two key residues depicted as red beads. The residues spending at least 75% of the trajectory inside these two regions are represented as blue beads. The Dijkstra's algorithm is used to compute the paths of minimal length on the graph (purple dashed lines) from the source to the sink region.

**Supplementary Table 1. Molecular dynamics simulations system setup.**

| | WT | $NgBR^{S249V}$ |
|---|---|---|
| Simulation box dimensions (Å) | 90.5 x 123.9 x 106.1 | 92.8 x 118.7 x 109.2 |
| No. of atoms | 102,113 | 112,615 |
| No. of water molecules | 33,973 | 34,505 |
| Salt concentration (KCl) | 0.15 M | 0.15 M |
